# A formula to estimate a researcher’s impact by prioritizing highly cited publications

**DOI:** 10.1101/058990

**Authors:** Aleksey V. Belikov, Vitaly V. Belikov

## Abstract

Here we suggest a new index to estimate the scientific impact of an individual researcher, namely the *S*-index. This index has been designed to emphasize highly cited, truly important works and to be minimally affected by poorly cited ones, without setting any arbitrary threshold. The first property makes it advantageous over the *h*-index, which does not discriminate between highly and moderately cited articles, while the second property - over the total number of citations, preventing the possibility of overclocking an index by publishing many trivial articles. Contrary to the *h*-index, which has an upper limit of the total number of publications regardless of their citation numbers, the S-index is not limited by the publication count. This allows scientists having few but very influential works to receive appropriate and respectable index values, which is impossible with the *h*-index. Moreover, only 10 most cited publications of an individual are typically required to calculate an *S*-index to 99% accuracy. Collectively, the *S*-index is principally different from the existing scientometric indicators and should facilitate better recognition of prominent researchers.

## Introduction

Scientific merits of researchers should ideally be evaluated by the impact of their works on particular fields of study and on scientific progress in general. To determine such impact is not easy, and sometimes decades pass before the true importance of a scientific discovery becomes clear. However, in many situations, such as during appointment of candidates to academic positions and distribution of research grants, it is required to make rapid and informed decisions based on somewhat limited data. Various quantitative indicators have been used for these purposes. However, existing bibliometric and scientometric indices are poorly suited to estimate scientific impact of an individual researcher. Below, we briefly describe their specifics and shortcomings.

*The total number of publications* is an indicator of productivity, but not impact. Not all publications make equal contributions to scientific progress. *The total number of citations* is a much more adequate indicator, because it reflects the influence that the publications of a scientist have had on the scientific community. However, a major disadvantage of this indicator arises from a potential contribution of a large number of poorly cited publications (and especially the contribution of self-citations). *The average number of citations per publication* reflects the impact of an average publication, but does not account for the total impact of the scientist’s work. This indicator is unreasonably high for scientists with few publications and unreasonably low for those with many. *The number of publications with “high” number of citations* suffers from the necessity to set an arbitrary threshold, above which the number of citations can be considered “high”. Moreover, this has the drawback of the first indicator discussed here in that quantity does not reflect quality. Two scientists with equal numbers of “important” publications can have considerable differences in the number of citations of those publications.

The *h-index* (Hirsch index) is defined as a maximal number of publications h, each of which has been cited at least h times^1^. Despite being quite easy to determine and truncating the tail of poorly cited publications, the h-index has a very significant drawback – it imposes the necessity of having a high amount of publications. For instance, a scientist who made a ground-breaking discovery, but published few articles, albeit with a very high number of citations, will have an *h*-index comparable to that of a starting postdoc. Gregor Mendel, the Father of Genetics (has 1 publication, see **Supplementary Table S4**), Isaac Newton, author of the Law of Universal Gravitation and the Laws of Motion (has 4 publications, see **Supplementary Table S1**), and Peter Higgs, predictor of the elementary mass particle and Nobel prize-winner (has 5 publications, see **Supplementary Table S10**) serve as real-world examples. These undeniably great scientists have h-indices of 1, 4 and 5, respectively. This obvious injustice is compensated by their deep public recognition and wide prominence of their discoveries; however, it does not add credibility to the *h*-index.

Another well-known scientometric indicator is the *g-index*. It is defined as the maximal number of the most-cited publications g, which have received *together* at least g^2^ citations^2^. In other words, each such publication should have *an average* of g citations. Although the *g*-index is better than the *h*-index at taking into account highly cited publications, it nevertheless suffers from the same drawback – a high *g*-index is not possible without a high number of publications.

## S-index principles and calculation

Having analysed the shortcomings of the existing bibliometric indicators in determination of a researcher’s scientific impact, we decided to develop an index that would be largely devoid of these drawbacks. Our main idea is that, during the calculation of an index of *impact*, it is necessary to increase the contribution of publications with a high number of citations and minimize the influence of poorly cited ones. This should prevent the possibility of overclocking an index by publishing many trivial articles and self-citing them. To prioritize publications with a high number of citations, we decided to use the squares of the number of citations for each publication, and calculate their sum. To prevent values of the index from becoming too large, and to contain them roughly within the range of the *h*-index, we decided to extract the root of the fourth degree from the resulting sum.

Thus, we propose a new index to estimate the scientific impact of an individual researcher, namely the ***S*-index** (from *square, sum* or *sigma*):

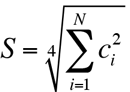

where *c_i_* – number of citations of the *i*-th publication, and *N* – total number of publications.

**Fig 1** shows citation data for the 50 most cited publications of 4 real, arbitrarily chosen scientists with different amounts and distributions of citations ( **A**, **B**, **C**, and **D**). The upper solid) curve on each graph reflects the distribution of the number of citations of each publication *c_i_*, plotted on the left *Y*-axis, according to the sequential number of publication *i* in the order of decreasing number of citations. Therefore, the area under this curve filled) equals the total number of citations *C*. The lower (dashed) curve reflects the distribution of the squared number of citations of each publication 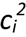 plotted on the right *Y*-axis, according to the number of publication *i*. Therefore, the area under this curve (hatched) equals the *S*-index to the fourth power (before extracting the root).

As can be seen from the figure, the contribution of publications to an S-index indeed progressively diminishes upon decrease in their citation numbers. It also can be noted that the S-index works equally well with anynumber and distribution of citations. Moreover, these examples demonstrate that, in most cases, 10 to 20 (rarely 30) publications are sufficient to calculate an S-index to high precision. This is possible due to the fact that scientific citations exhibit the behaviour of *preferential attachment* which results in their distributions according to power law^3^.

**Fig. 1.**
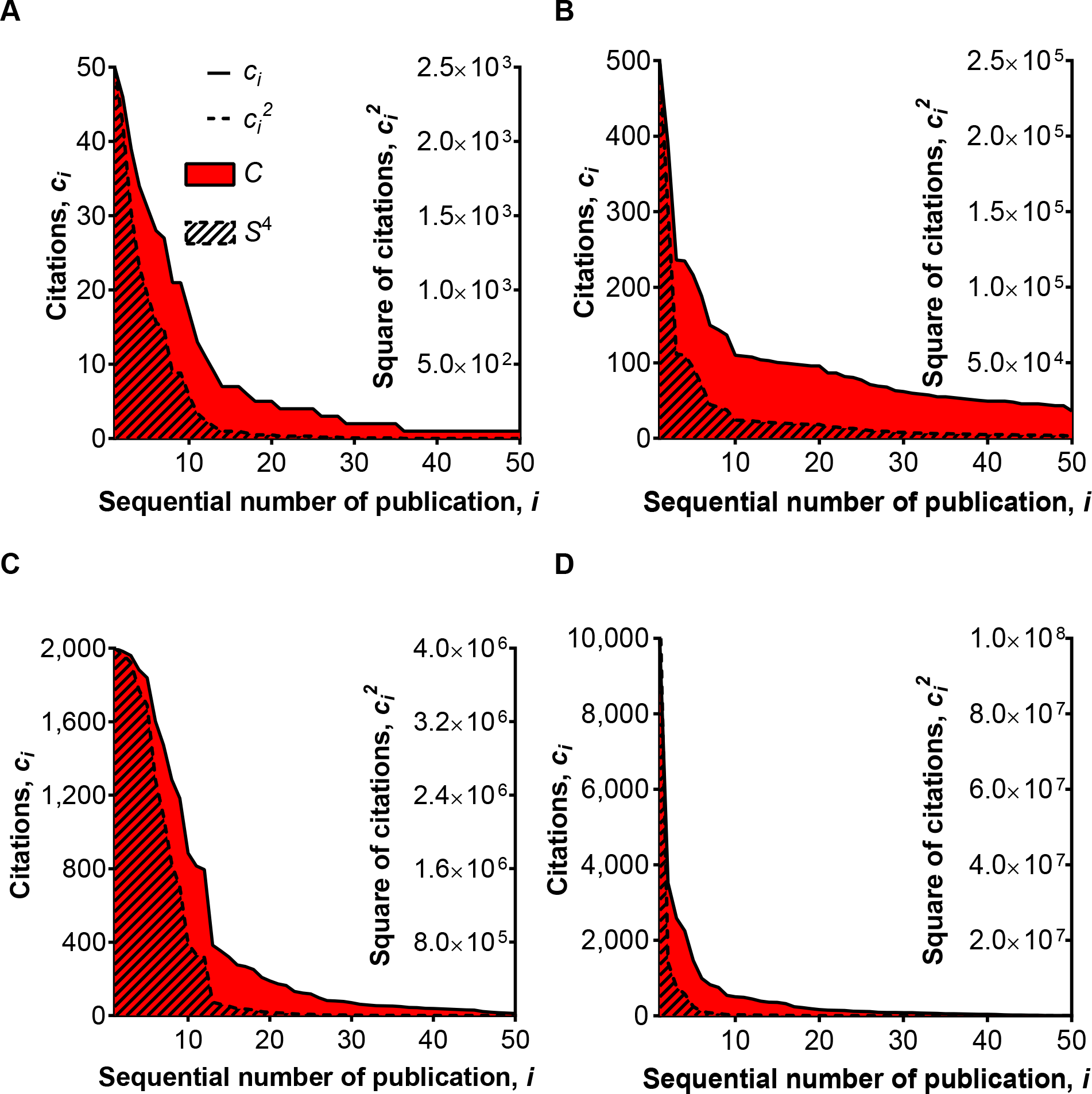
Relative contribution of individual publications to the total numbers of citations and to *S*-indices. Examples from 4 actual scientists (**A**, **B**, **C**, **D**) are shown. Publications are sorted in the order of decreasing number of citations. *C* - the total number of citations, *S* – an *S*-index. Legend in **A** applies also to **B**, **C**, and **D**.

As an example, we will now calculate an *S*-index for Isaac Newton, who has only 4 scientific publications (see **Supplementary Table S1** for publication list and citation data):

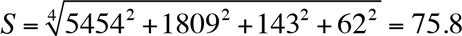

As can be seen, the difference with a Hirsch index *h*= 4 is quite considerable.

**Table 1.** Comparison of bibliometric indicators for influential scientists.

| Scientist | *N* | *C* | *C/N* | *h* | *g* | *S*<sub>10</sub> | *S*<sub>20</sub> | *S*<sub>30</sub> |
| --- | --- | --- | --- | --- | --- | --- | --- | --- |
| Isaac Newton | 4 | 7468 | 1867 | 4 | 4 | 75.8 | 75.8 | 75.8 |
| Michael Faraday | 23 | 6288 | 273 | 18 | 23 | 55.5 | 55.5 | 55.5 |
| Charles Darwin | 29 | 84575 | 2916 | 27 | 29 | 207.8 | 207.9 | 207.9 |
| Gregor Mendel | 1 | 2442 | 2442 | 1 | 1 | 49.4 | 49.4 | 49.4 |
| Max Planck | 32 | 6787 | 212 | 27 | 32 | 44.4 | 45.0 | 45.0 |
| Albert Einstein | 130 | 72286 | 556 | 74 | 130 | 133.0 | 134.7 | 135.0 |
| Niels Bohr | 52 | 21069 | 405 | 38 | 52 | 71.0 | 72.2 | 72.3 |
| Louis de Broglie | 70 | 6482 | 93 | 40 | 70 | 38.4 | 38.9 | 39.1 |
| Francis Crick | 65 | 43758 | 673 | 54 | 65 | 110.2 | 111.6 | 111.9 |
| Peter Higgs | 5 | 11903 | 2381 | 5 | 5 | 82.5 | 82.5 | 82.5 |
| **Mean** | **41** | **26306** | **1182** | **29** | **41** | **86.80** | **87.35** | **87.44** |
| **%RSD** | **97** | **114** | **92** | **82** | **97** | **60** | **60** | **60** |

*N* – the total number of publications, *C* – the total number of citations, *C/N* – the average number of citations per publication, *h* – an *h*-index, *g* – a *g*-index, *S* – an *S*-index, %RSD – a relative standard deviation. *S*-indices were calculated based on 10 (*S_10_*), 20 (*S_20_*) and 30 (*S_30_*) most cited publications.

Meanwhile, a hypothetical postdoc who also has 4 publications and *h* = 4 but far fewer citations, achieves an *S*-index that is comparable with *h*:

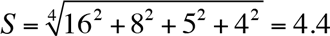

Thus, the sensitivity of the *S*-index considerably exceeds that of the h-index. It should be noted that, in contrast to an *h*-index, an *S*-index is not an integer, and can be calculated to any desired precision.

## S-index validation and analysis

**Table 1** presents various bibliometric indicators obtained for 10 arbitrarily selected influential scientists, and calculated using citation data from Google Scholar ^a^ see **Supplementary Tables S1-S10** for the citation data used in this work).

There are several interesting observations that can be made from these data. First, there is a notably large scatter in the values of presented indicators for the different scientists. This is reflected in the high values of relative standard deviation (%RSD). It is logical to suppose that the lower is the scatter in the values of a parameter for the members of some set, the better this parameter reflects commonality of those members. In the particular case of estimating scientific impact, a more successful indicator will have a lower scatter for the selection of influential scientists, i.e., will have higher predictive power. As can be seen from **Table 1**, the lowest scatter (%RSD) is exhibited by *S*-indices.

As mentioned above, the *h*-index is not applicable to scientists with a low number of publications *N*, because *N* acts as a harsh limiting factor for h. This can be clearly seen in **Fig 2A**, where the correlation between *h* and *N* for scientists from **Table 1** is very high (R^2^ = 0.977). As a result, almost a third of world-famous scientists from our list have unfairly low *h* (≤ 5). Meanwhile, *S*-index values for the same scientists do not correlate with the publication numbers (R^2^ = 0.115). However, even more interesting is the mean *h*-index value for the influential scientists, which equals 29. Such an *h* value is not considered exceptional. However, the mean *S*-index value for the same scientists equals 87.

**Fig. 2.**
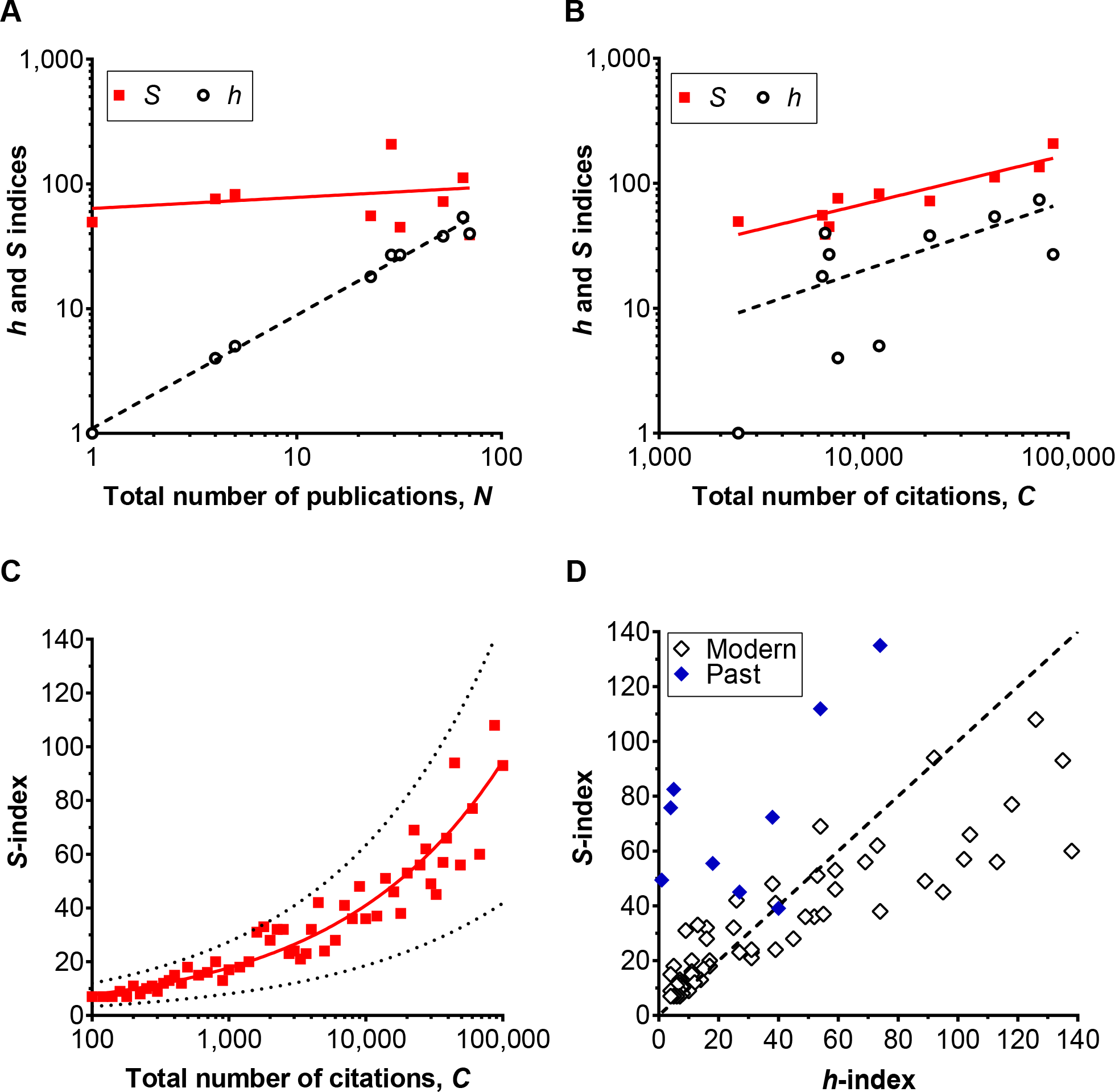
Correlations of *S*- and *h*-indices with the total numbers of publications, the total numbers of citations, and with each other. (**A** and **B**) Correlations of *S*- (solid squares) and *h*- (open circles) indices with the total numbers of publications **A**) and the total numbers of citations **B**), for scientists from Table 1. Solid line – log-log regression for *S*, dashed line – log-log regression for *h*. (**C**) Correlation of *S*-indices with the total numbers of citations, for contemporary scientists. Solid line – log-log regression, dashed lines – 99% prediction band. (**D**) Comparison of *S*-indices with *h*-indices, for contemporary scientists open diamonds) and scientists from Table 1. solid diamonds). Dashed line: *h*=*S*. In all panels, each dot represents an individual scientist.

**Fig 2B** shows that, for scientists from Table 1, *h*-index values do not correlate with the total numbers of citations *C* (R^2^ = 0.116), a much more important indicator of scientific impact than the total number of publications. By comparison, the *S*-index values demonstrate a strong correlation with *C* (R^2^ = 0.855). All the foregoing indicates that the *S*-index is better suited for estimation of scientific impact, whereas the *h*-index is more an indicator of productivity.

We demonstrated that the *S*-index, unlike the *h*-index, gives the proper credit to the exceptional world-famous scientists from the past. It is interesting, however, how the *S*-index behaves on contemporary scientists. To answer this question, we have randomly selected Google Scholar profiles^b^ of about 60 scientists with total citation counts ranging from 100 to 100000. As can be seen from **Fig 2C**, *S*-index values for those scientists correlate strongly with the total numbers of citations *C* (R^2^ = 0. 921). Nevertheless, the scatter is large enough to not allow for substitution of the *S*-index by *C*.

In **Fig 2D**, open diamonds show the direct comparison between *S*- and *h*-indices for contemporary scientists. It can be noted that the *S*-index typically has equal or higher than *h* values for scientists with low *h*-indices (*h*<20), comparable values for those with medium *h*-indices 20<*h*<60), and lower than *h* values for individuals with high *h*-indices (*h*>60). This is contrasting with the influential scientists from the past (solid diamonds), who have *S*-indices considerably higher than *h* in all cases but one. In other words, for each value of *S* impact) modern scientists have larger *h* (productivity). This distribution most likely reflects the current trend to publish more articles, not necessary increasing the cumulative impact of one’s research.

The *g*-index also demonstrates an interesting property. As can be seen from **Table 1**, for the influential scientists considered, *g-* indices turn out to be identical to publication numbers *N.* In these cases, *N* behaves as a limiting factor, because the total number of citations *C* for influential scientists is always high. Consequently, the g-index in its classical definition is not applicable for evaluation of scientists with *C* ≥ *N*^2^. In note added in proof, Egghe^2^ suggests to add “fictitious articles with 0 citations” to overcome this problem. However, this rather counterintuitive procedure reduces the g-index simply to the square root of the total number of citations, rounded down to the nearest integer.

We will now analyse the difference between *S*-indices that were calculated based on 10, 20, and 30 most-cited publications (see **Table 1**). Here, the mean values of Si_0_ and S_20_ constitute respectively 99.3% and 99.9% of S_30_. Thus, when citation data for all publications are unavailable and/or when an *S*-index is calculated manually, it is acceptable to use citation data concerning 20 most-cited publications. For quick, rough estimation, only 10 most-cited publications are required.

## S-index modifications

Sometimes it is required to estimate the impact of only *recent* publications of a researcher, in order to assess the scientist’s ongoing research achievement. For such cases, we suggest a modification of the *S*-index, which we call the *5-year S*-index (^5yr^*S*). This index is calculated in the same way as a standard *S*-index, except that outputs published solely during the last 5 years are used.

Another interesting question that may arise is how relevant are *all* publications of a scientist *at present*. For this purpose, we suggest the *relevance S*-index (^rlc^*S*), which is calculated in the same way as a standard *S*-index, except that *citations* made only during the last 5 years for *all* publications of a scientist are utilized.

If a *threshold-free* solution is desired to compensate for aging of publications, we suggest the *age-normalized S*-index (^age^*S*), which is calculated in the same way as a standard *S*-index, except that for each publication the number of citations is divided by the age of the publication in years.

A common drawback of most widely adopted indicators of scientific influence is their total insensitivity to the contribution of individual scientists (as co-authors) to a given publication. For example, a scientist who only briefly participated in the creation of an article with 30 co-authors is allocated the same contribution to the indicators as another scientist who performed the bulk of the work in a two-author article. To eliminate this flaw, we suggest the *contribution-compensated S*-index (^cnb^*S*), which is calculated in the same way as the standard *S*-index, except that for each publication the number of citations is divided by a coefficient reflecting the contribution of the scientist to the publication. The total number of co-authors could serve as a straightforward candidate for this coefficient. However, more sophisticated inferring of the *contribution coefficient* from the list of coauthors is intimately linked to how the authorship distribution has been arranged, which is often field-specific^4^, and therefore is a non-trivial task. A standardized electronic system to register co-author contributions that is integrated with manuscript submission systems and ORCID^c^ would be of great help. Such a system is currently being tested^5^. Alternatively, an automated computational way to determine an author’s credit share can be employed^6^.

In conclusion, we have proposed a new index to estimate the scientific impact of an individual researcher and demonstrated its advantages over the existing bibliometric indicators. We hope that its practical application will aid a better recognition of truly influential scientists.

## Acknowledgements

The authors are grateful to Prof. Alistair Borthwick, Dr. Alexander Barmashov and Vladimir Hrubilov for critical reading of the manuscript and useful comments.

## Author contribution

A.V.B. co-created the *S*-index formula, collected and analysed the data, interpreted the results and wrote the manuscript. V.V.B. cocreated the *S*-index formula, interpreted the results and revised the manuscript. All authors gave final approval for publication.

## Footnotes

a https://scholar.google.com

b https://scholar.google.com/citations?mauthors=&view_op=search_authors

c http://orcid.org

